# Development of a Low-cost Extrusion Multi-nozzle 3D Bioprinter for Complex Constructs

**DOI:** 10.64898/2026.10.14.756591

**Authors:** Kinam Hyun, Annika Müller, Nima Tabatabaei Rezaei, Youssef Fahmy, Jaemyung Shin, Chaenyung Cha, Keekyoung Kim

**Affiliations:** Department of Mechanical and Manufacturing Engineering, Schulich School of Engineering, University of Calgary, Calgary, AB, T2N1N4, Canada; Helmholtz Institute for Translational Oncology Mainz (HI-TRON Mainz), 55131 Mainz, Germany; Division of Stromal Immunology, German Cancer Research Centre (DKFZ), 69120 Heidelberg, Germany; Department of Medicine, Cumming School of Medicine, University of Calgary, Calgary, AB T2N 1N4, Canada; Snyder Institute for Chronic Diseases, University of Calgary, Calgary, AB T2N 1N4, Canada; Department of Biomedical Engineering, Johns Hopkins University, Baltimore, MD 21205, USA; Department of Biomedical Engineering, Schulich School of Engineering, University of Calgary, Calgary, AB, T2N1N4, Canada; Department of Materials Science and Engineering, Ulsan National Institute of Science and Technology, Ulsan 44919, Republic of Korea

**Author notes:** Corresponding author: Dr. Keekyoung Kim.

**Keywords:** Additive manufacturing, Multi-material bioprinting, Extrusion-based bioprinting, Bioprinter calibration

## Abstract

Extrusion-based multi-material bioprinting is a promising approach for fabricating complex tissue constructs, but current systems often require expensive hardware or support only one or a few numbers of bioinks. In this study, a low-cost, compact, and versatile multi-nozzle bioprinter (MNB) system was developed for pneumatic extrusion of up to 10 different inks using a rotational head. The system was integrated with a commercial 3D printer platform and controlled using custom G-code commands. Backlash compensation and nozzle-specific calibrations were experimentally determined to minimize positioning errors and airflow misalignment. The developed system showed stable repeatability in both clockwise and counterclockwise movements. Lines and various structures were successfully printed with different nozzle-switching directions and printing sequences. Representative structures including a diamond pattern, a multilayer scaffold, a pyramid structure, and a kidney-like construct were successfully fabricated. These results demonstrate that the proposed MNB system can produce complex 2D and 3D multi-material structures with good positional accuracy and repeatability.

## 1. Introduction

The need for advanced tissue engineering arises from the demand for a better understanding of tissue behaviors ^[1]^, improved drug development ^[2–4]^, and artificial organ fabrication ^[5]^. However, replicating the complexity of native tissue structures remains challenging. Organoid techniques have therefore been widely utilized because they can relatively easily form tissue-like structures through self-organizing cell aggregates. Although organoids can mimic certain aspects of native tissues, they are commonly limited to micrometer to millimeter-scale sizes ^[6–8]^. They also face challenges in reproducibility, long-term culture maturity and the replication of complex microenvironments such as microvasculature, which limits their scalability toward larger tissue and artificial organ fabrication. 3D bioprinting has therefore gained attention as a promising approach because it enables precise spatial control and scalable fabrication of complex tissue constructs ^[9,10]^. For example, extrusion bioprinting has been applied to high-throughput organoid production ^[11,12]^, while stereolithography (SLA) printing has been used for microarchitectural patterning ^[13]^.

Among various bioprinting methods, extrusion-based bioprinting is widely used because of its material versatility, fast printing speed, and high cell viability. Nevertheless, extrusion-based bioprinting still has several limitations. These include complex calibration processes ^[14,15]^, relatively low printing resolution ^[16]^, the use of only one or two bioinks ^[10,14]^, and the high cost of commercial bioprinters ^[17,18]^. For example, vascular structures often require hollow channels that are generated using sacrificial materials ^[19]^. To overcome these limitations, extrusion-based multi-material bioprinters have recently emerged ^[20–24]^. These systems can be classified into single-nozzle and multi-nozzle approaches. Single-nozzle systems generally have mechanically simpler hardware and easier alignment because only one dispensing position is used. However, they often require more complex fluidic design to control flow rate, viscosity, shear stress, and rheological compatibility between different bioinks. Examples of single-nozzle systems include bundled microfiber nozzles ^[25–27]^, preset cartridge systems ^[9]^, and mixing nozzles ^[28]^. In contrast, multi-nozzle systems use multiple nozzles to dispense different bioinks independently ^[18,29–32]^. This approach provides more flexibility in bioink selection and independent material control because each bioink can be individually controlled with minimal cross-contamination and without requiring complex fluidic compatibility between materials. These advantages provide opportunities for fabricating more complex and application-specific tissue constructs. For example, Sebastien G. M. Uzel et al. developed a bioprinter with an adjustable nozzle height to print on complex substrate topographies such as skin ^[33]^.

In this paper, we propose a multi-nozzle 3D bioprinter (MNB) system that can operate with 10 different bioinks. This system offers several advantages including cost-effectiveness, simple operation, and versatile control. These advantages are achieved by installing a rotational head on a commercial 3D printer and controlling it with G-code generated by open-source software. The MNB system has potential for printing cell-laden bioinks and can be operated inside a biosafety cabinet.

## 2. Materials and Methods

### 2.1 3D modeling and fabrication of MNB system

3D models were designed using 3D CAD software (Autodesk Inventor Professional 2024, Autodesk, USA) and exported as STL files for 3D printing. The body head and adapters were printed using resin (Clear Resin V4, Formlabs, USA) with an SLA 3D printer (Form 3+, Formlabs, USA). Other components including the body and rotational head were printed using PLA+ filament with a fused deposition modeling (FDM) 3D printer (Ender3 Pro, Creality, China). All print designs were sliced and converted into G-code using corresponding printer software.

### 2.2 MNB system configuration

The MNB system was designed to pneumatically extrude 10 different inks using the developed rotational head (Fig. 1 and Fig. S1). As shown in Fig. 1C, it mainly consists of a dispensing system, a 3D printing station, and the rotational head module. The system was designed to be compatible with commercial 3D printers and consumables. It was also developed based on a previously reported in-house 3D bioprinter ^[34,35]^. Briefly, the existing filament printhead on a commercial 3D printer (Ender3 Pro, Creality, China) was disassembled, and the nozzle cooling fan wires were reconnected to a digital pneumatic dispenser (ML-5000XII, Musashi Engineering Inc., Japan) through the developed PCB circuit. The PCB contains voltage regulators that properly transmit signals to the dispenser and enable G-code control (Table 1).

**Table 1.**
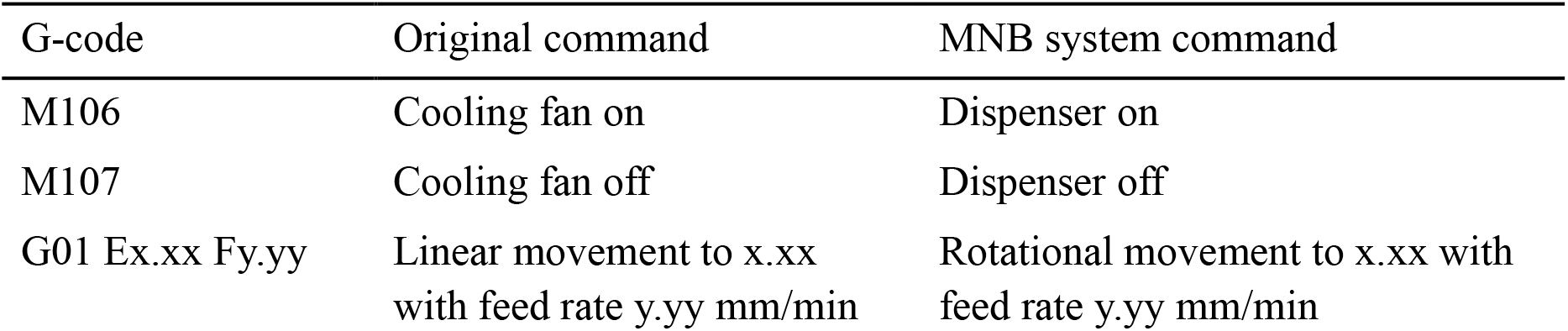
Assigned G-codes for MNB system.

**Fig. 1.**
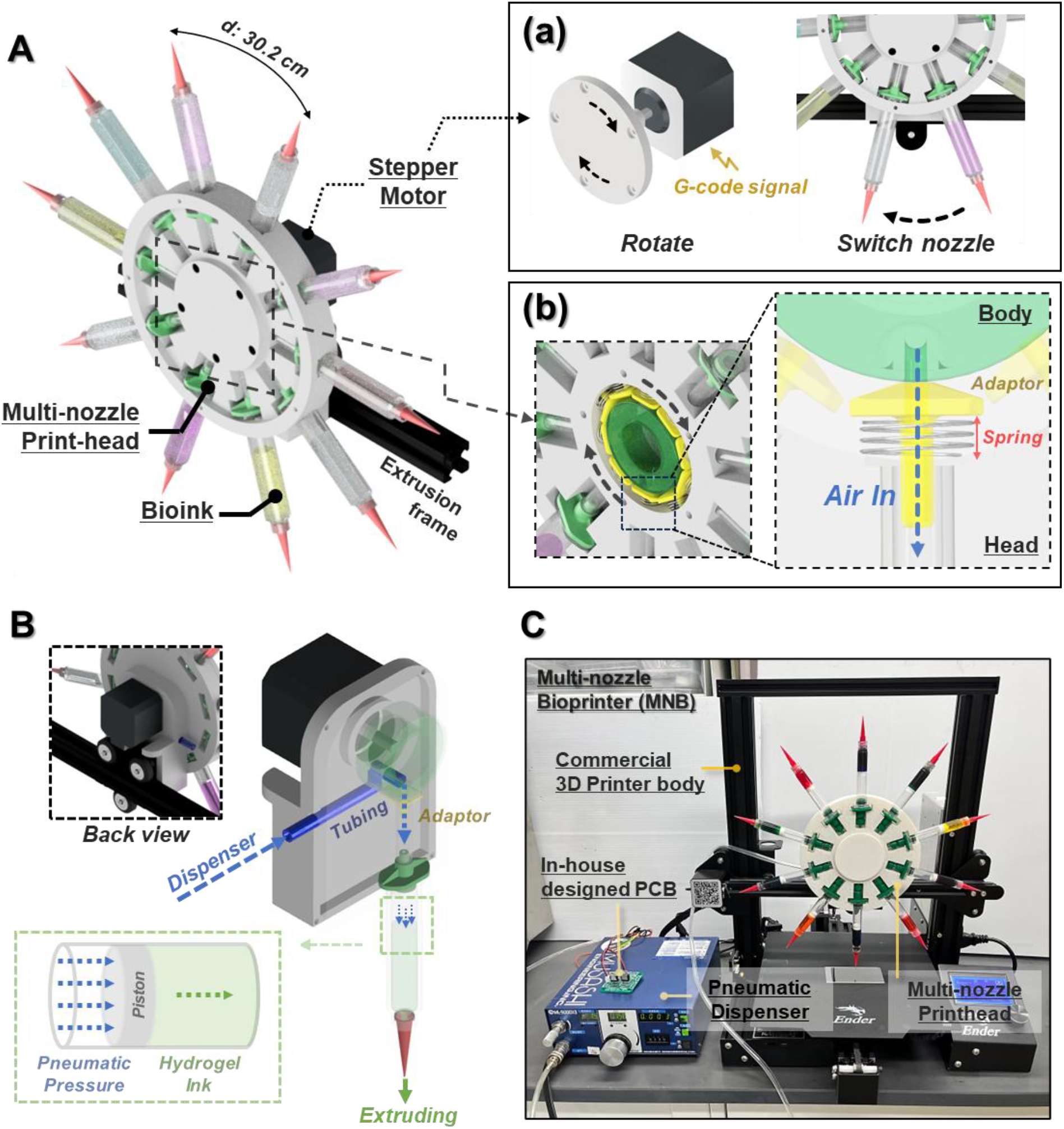
Overview of the 3D multi-nozzle bioprinter (MNB) system. A) Design of the rotational head with a stepper motor mounted on a commercial 3D printer. (a) Mechanism of nozzle switching by controlling the motor. (b) Internal structure of the rotational head including spring adapters for air flow. B) Schematic of the pneumatic flow path within the system. C) Photograph of the complete MNB system including the rotational head, commercial 3D printer body, in-house developed PCB, and a pneumatic dispenser.

The rotational head module mainly consists of a rotational head plate, a main body frame, and a stepper motor (Fig. 1A). The rotational head plate measured 140 mm × 14 mm, and 302 mm × 17 mm including the nozzle tips, and had 10 slots for syringe cartridges and adapters. 3 mL plastic syringe cartridges with pistons, and various needle tips were purchased from CELLINK (CA, USA), while syringe adapters were purchased from Amazon (3 mL glue adhesive dispenser syringes connector adapter, USA). The wings and ends of the syringe adapters were cut to fit into the slots of the head. 3D-printed air-sealing adapters with 8 mm diameter springs were inserted at the center of the head plate. These spring adapters help maintain the head plate at the center and prevent air leakage between the head plate and the body frame (Fig. 1A (b)). After inserting ¼-inch silicone tubing, the main body frame was assembled with glue and screws. This tubing was connected to the pneumatic dispenser and delivered air to the piston in the cartridge (Fig. 1B). The assembled body frame was then mounted on the 3D printer bracket, and the rotational head plate was then placed on the rotational shaft. A stepper motor (Nema 17, 45N·cm, StepperOnline, USA) was attached to the back of the body frame, and the cable from the filament feeding motor was connected. This connection enables G-code control of the rotational head (Table 1). The stepper motor was connected to the rotational head through the motor coupling bracket which was then fixed to the rotational head plate with screws (Fig. 1A (a)).

### 2.3 Ink preparation

Inks for MNB system characterization were prepared using Pluronic F-127 (PF-127, Sigma-Aldrich, USA). Pluronic powder was dissolved at 40% (w/v) in cold distilled water. The solution was repeatedly vortexed and stored at 4 °C until the powder was completely dissolved. After dissolution, the solution was colored with different food dyes and kept at 4 °C until further use. Before printing, the prepared inks were loaded into capped cartridges, and care was taken to ensure that no air bubbles were present in the cartridges. The inks were then kept at room temperature for at least 1 hour to stabilize their rheological properties.

### 2.4 G-code generation and 3D extrusion bioprinting

The G-codes for the 3D-printed constructs were designed and generated using FullControl GCODE Designer ^[36]^. FullControl GCODE Designer is open-source software that enables direct generation of G-code by explicitly defining print paths and printing parameters without requiring STL conversion or conventional slicing software (Fig. S2). This approach also allows incorporation of custom G-code commands, such as nozzle switching and pneumatic dispensing control, which are particularly useful for the MNB system. All MNB system operations and G-code file uploading were conducted using the open-source 3D printing host software Pronterface (Printrun 2.2.0, GNU General Public License, version 3).

### 2.5 Image processing and analysis

A microscope (Revolve, Echo, USA) was used to measure the width of printed lines. G-code paths were visualized and captured using NC viewer (https://ncviewer.com) ^[37]^. A USB digital camera (Jiusion, Shenzhen, China) and its software (Xploview, Jiusion, Shenzhen, China) were used to record videos and capture images for calibration. Images extracted from the videos were analyzed and processed using ImageJ. Accuracy and repeatability were evaluated using position error, which was calculated using Equation (1):

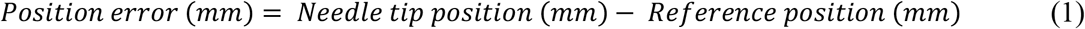

The needle tip position was measured by marking the tip along the x-axis using a 10 mm reference scale, which was manually defined by drawing a line on a ruler placed behind the tip. The reference position was defined as the initial needle tip position along the x-axis.

### 2.6 Statistical analysis

All data are presented as mean values accompanied by the standard deviation. Statistical analysis was conducted using Prism 10 (GraphPad, USA). To determine statistical significance, a one-way analysis of variance (ANOVA) was employed along with Tukey’s multiple comparison test. Differences were regarded as statistically significant when the p-value < 0.05, and the significance levels were indicated as follows: ns (not significant) for p > 0.05, *p < 0.05, and **p < 0.01.

## 3 Results

### 3.1 Calibration of MNB system

To minimize position error and airflow misalignment during ink extrusion, it is necessary to determine the optimal rotational parameters for nozzle switching. Key rotational parameters including backlash, rotational speed, and rotational increments between each nozzle were therefore evaluated. All calibration tests were performed within ±E0.005 for the rotational E-value and ±1 mm for nozzle position accuracy. Cylindrical needle tips with a gauge size of 25G (ID: 0.26 mm) were used for the calibration process. First, the backlash of the stepper motor was measured during single-nozzle movement following forward and backward rotation. Because the head was rotated using a stepper motor, inherent backlash was present in the system. As shown in Fig. 2A, the nozzle position was displaced by approximately 26 mm without backlash compensation. Therefore, a backlash compensation value of E1.025 was experimentally determined and applied to the rotational movement.

**Fig. 2.**
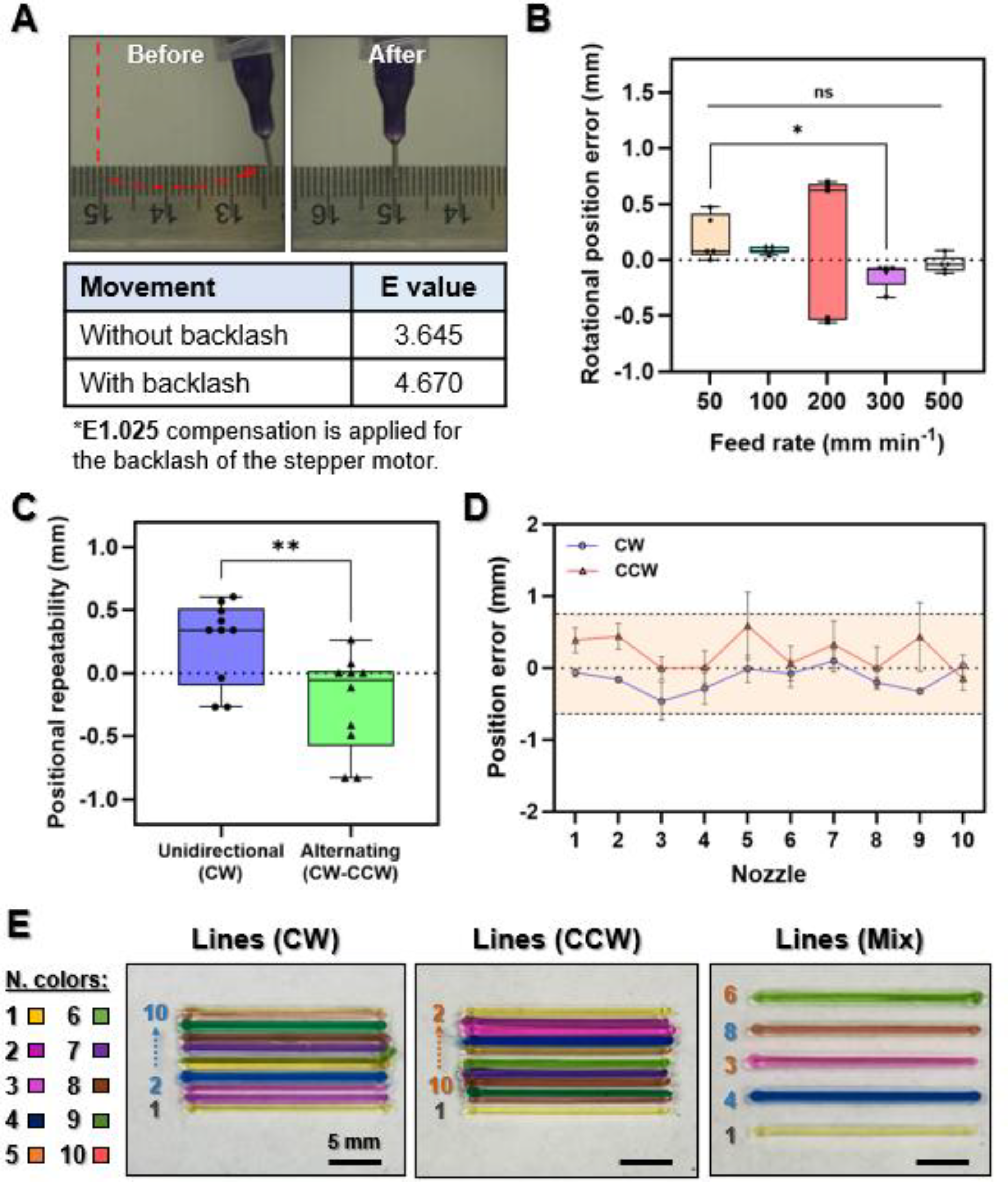
MNB system characterization. A) Nozzle tip positions under preloaded and non-preloaded backlash conditions. B) Rotational calibration at different motor feed rates (n = 5), C) Positional repeatability of the rotational head under different movement strategies (n = 10). D) Accuracy of nozzle positioning for each nozzle (n = 4). E) Line printing results under different rotational directions using 40% (w/v) PF-127.

The graph in Fig. 2B illustrates the position error as a function of the rotational feed rate. To determine the appropriate rotational speed of the head, the head plate was fully loaded with 10 inks and tested. The head was rotated through 360° in the clockwise (CW) direction, and this process was repeated 5 times. At 50 mm/min, the larger position error may be attributed to static friction, and stuttering movement was observed during rotation. The largest error was observed at 200 mm/min. This is likely because E37.795 was selected for one full rotation, and a feed rate of 200 mm/min could not evenly divide the movement interval. As a result, the errors fluctuated before and after the zero position. At feed rates above 300 mm/min, the effects of static friction became negligible, resulting in much smaller position errors. However, in the single-nozzle movement experiment, the error increased substantially. This may be due to the short rotational increment, which increases the influence of transient response and mechanical instability. Therefore, 100 mm/min (F100) was selected as the nozzle-switching speed (E-axis feed rate) for the remaining experiments.

To demonstrate the repeatability of the rotational head, the head was rotated through 360° either only in the CW direction or in alternating CW and counterclockwise (CCW) directions. These two conditions represent movements without backlash and with backlash, respectively. This process was repeated 10 times. One full rotation was experimentally determined to correspond to an E-value of 37.795. As shown in Fig. 2C, the average position error of unidirectional movement was 0.255 ± 0.326 mm, whereas alternating movement showed an average position error of - 0.234 ± 0.385 mm. The difference in error direction and the larger variation observed during alternating movement may be attributed to backlash introduced during directional changes. Although the two movement conditions showed significantly different errors, both were within the desired accuracy range of ±1 mm.

Single-nozzle positioning was also evaluated. However, each nozzle required individual coordinate calibration due to variations in components. The plastic syringe cartridges were inherently and randomly bent, likely because of the injection molding process during manufacturing (Fig. S4). Even slight bending caused millimeter-scale displacement and pressure variation, which degraded the overall performance of the MNB system. In addition, minor rotational friction and elastic compliance from the 3D-printed components could cause rotational displacement. Therefore, each nozzle was assigned to its own position, and manual calibration was performed. The Y-axis was calibrated by measuring the tip position on the millimeter grid, whereas the X and Z-axes were calibrated using a ruler placed behind the tip. As shown in Fig. S5, the most severely bent cartridge exhibited a displacement of 3.5 mm. Meanwhile, the X-axis was calibrated by adjusting the E-value after eliminating backlash and was performed within ±1 mm (E0.005). The E-values for each nozzle in both CW and CCW directions were experimentally determined and listed in Table S1. Although no significant trend was observed in the E-values, the range of variation was approximately 4.27% for CW movement and 3.85% for CCW movement, while the overall error was approximately 0.98% and 0.96%, respectively. These values were sufficiently small to allow stable operation of the system. After manual calibration, the positioning accuracy of each nozzle was evaluated (Supplementary Video S1). The trend of position errors was slightly shifted toward the negative side for CW rotation and the positive side for CCW rotation. This may be because the calibration resolution was limited to E0.005. However, the overall standard deviation was below 0.5 mm, and the range of positioning error was within ±0.589 mm. This indicates that the positioning accuracy was sufficient for stable nozzle switching and ink extrusion.

### 3.2 Printing performance of the MNB system

To demonstrate the printing performance of the MNB system, lines were printed using 10 nozzles with different ink colors. Cylindrical needle tips with a gauge size of 25G (ID: 0.26 mm) and a printing speed of 300 mm/min were used for line printing. First, the nozzle was moved to the calibrated position using absolute coordinates, whereas all subsequent movements were conducted using relative coordinates. Although nozzle positions were calibrated, pressure variation between nozzles remained. Therefore, the pressure was adjusted while a 10 mm pre-extruded line was printed at the side to stabilize the extrusion before actual printing. After printing each line, the head was rotated using the designated E-value and moved to the calibrated position of the next nozzle. This process was repeated until the entire sequence was completed. As shown in Fig. 2E and Supplementary Video S2, 20 mm lines with 1 mm spacing were successfully printed using different nozzle-switching directions. The printed lines were well aligned in both CW and CCW movements, indicating stable nozzle positioning and repeatable switching performance. In addition, the mixed nozzle-switching sequence also showed accurate nozzle selection without noticeable misalignment between lines. This result demonstrates that the MNB system can maintain consistent printing quality regardless of nozzle-switching direction or printing order.

### 3.3 Printing of various structures

To further demonstrate the versatility of the MNB system, several structures with different geometries and printing strategies were fabricated. Fig. 3A illustrates the overall printing workflow of the MNB system, including printing, moving to the next layer, and nozzle switching. During multilayer printing, the head rotated to select the designated nozzle after each layer was completed. The overall printing workflow from design generation to printing is shown in Fig. S6. A 10 × 10 mm diamond structure was successfully printed using 10 nozzles with different-colored ink (insert), as shown in Fig. 3B. The structure showed clear separation between colors and accurate positioning of each nozzle.

**Fig. 3.**
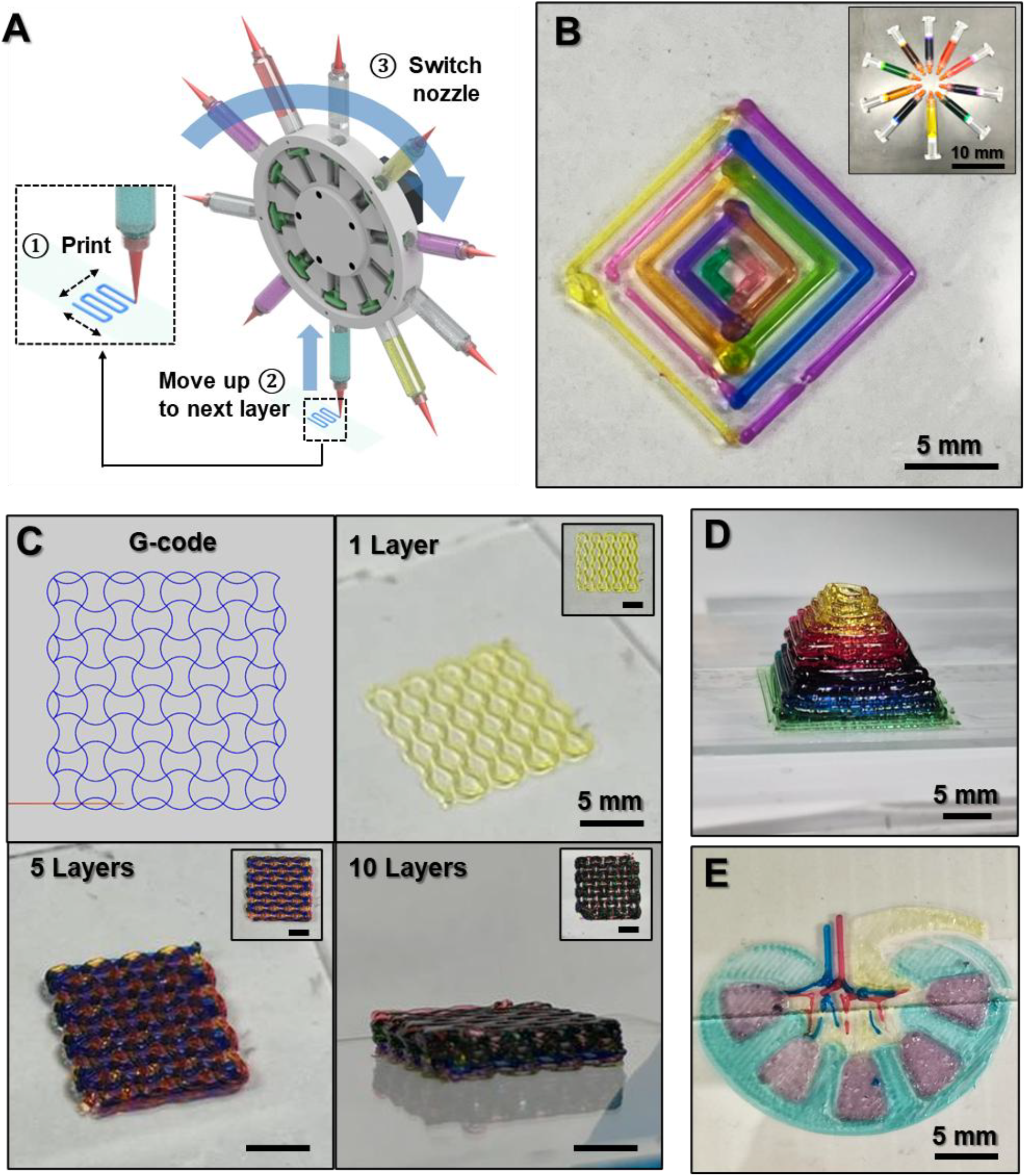
Demonstration of 3D constructs using the MNB system. A) Overview of the overall printing process. B) Diamond pattern (10 mm x 10 mm) printed using ten nozzles, C) Printing of 10-layer auxetic pattern scaffold (16 mm x 16 mm x 4 mm) including G-code path and images after 1, 5, and 10 layers printed (scale bars = 5 mm). D) Pyramid structure (20 mm x 20 mm x 15 mm). E) Kidney-like structure using 5 different nozzles (30 mm x 21 mm).

Fig. 3C shows the printing of a 10-layer auxetic scaffold with dimensions of 16 mm × 16 mm × 4 mm. The corresponding G-code path is shown on the top left. Two different wavy patterns were repeatedly stacked layer-by-layer at a 90° orientation. Images after 1, 5, and 10 printed layers demonstrate that the scaffold geometry was maintained throughout the multilayer process (Supplementary Video S3). In particular, the periodic pattern was preserved without significant collapse or distortion as the number of layers increased. A pyramid structure was also fabricated, as shown in Fig. 3D. The structure was printed by gradually reducing the printing area at each layer, resulting in a stable 3D geometry with distinct color transitions between layers. Finally, a kidney-like structure was printed, as shown in Fig. 3E. Different colors were used to represent individual regions of the structure, demonstrating that the MNB system can selectively deposit multiple inks within a single construct. These results indicate that the MNB system can fabricate not only simple line patterns but also complex 2D and 3D structures with multiple inks.

## 4. Conclusion

In this study, a MNB system equipped with a rotational head was successfully developed and characterized. The developed system was capable of pneumatically extruding 10 different inks using a single rotational head integrated with a commercial 3D printer platform. The rotational head enabled stable nozzle switching through G-code control, and key parameters including backlash, rotational speed, and nozzle-specific calibration were systematically optimized.

The results demonstrated that the MNB system could achieve stable nozzle positioning and repeatable switching performance in both CW and CCW directions. After calibration, the system showed sufficiently small positioning errors for reliable operation. The printing performance was validated through aligned line printing using 10 different nozzles, as well as through fabrication of various 2D and 3D structures including diamond patterns, auxetic scaffolds, pyramid structures, and kidney-like constructs. These results indicate that the proposed MNB system can fabricate complex multi-ink structures with good positional accuracy and repeatability.

However, several limitations remain in the current system. Pressure differences depending on ink viscosity and nozzle diameter still require additional adjustment before printing. In addition, manual leveling of the Z-axis and individual nozzle calibration are necessary prior to printing because of component variation and cartridge deformation. Future improvements in cartridge quality, such as replacing plastic cartridges with glass cartridges or higher-precision components, may reduce calibration requirements. Furthermore, image-based monitoring combined with machine learning analysis has the potential to enable automatic calibration and process optimization in future systems ^[35]^.

The proposed rotational head also has potential for integration with additional functional modules, such as light sources for photocrosslinking, biosensing electrode tips, and magnetic cell mixers. In the future, the MNB system could be further expanded toward organ-on-a-chip applications ^[38]^ and advanced biofabrication using more sensitive cell sources such as induced pluripotent stem cells (iPSCs), which require improved cell viability and printing conditions ^[39]^.

## Acknowledgement

This work was supported by a Natural Sciences and Engineering Research Council of Canada (NSERC) Discovery Grant (RGPIN-2014-04010) and Canada Foundation for Innovation John R. Evans Leaders Opportunity Fund.

