## Supplementary Information for "Development of a Low-cost Extrusion Multi-nozzle 3D Bioprinter for Complex Constructs"

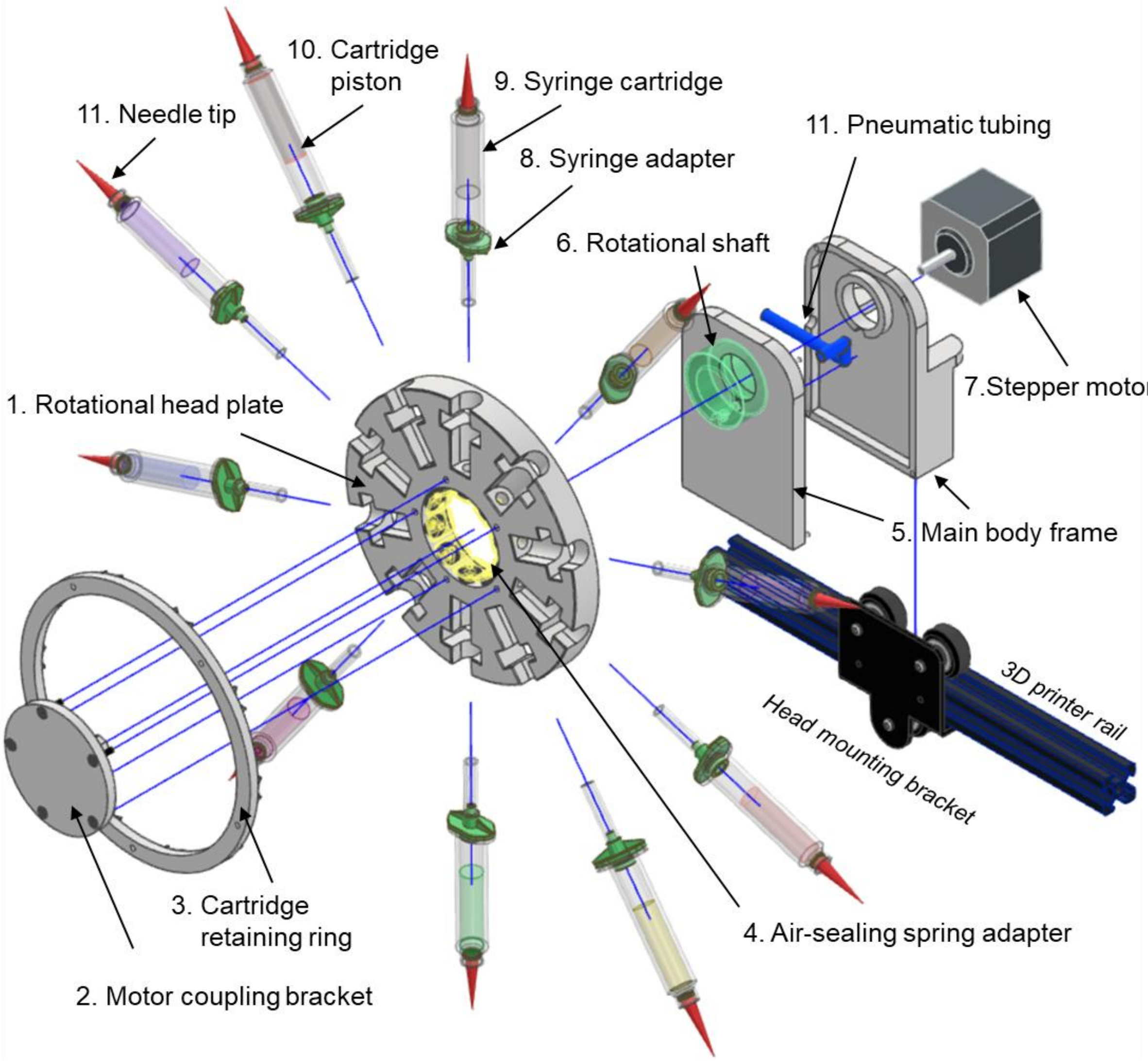


**Fig. S1**. Exploded view of the multi-nozzle bioprinter (MNB) system.

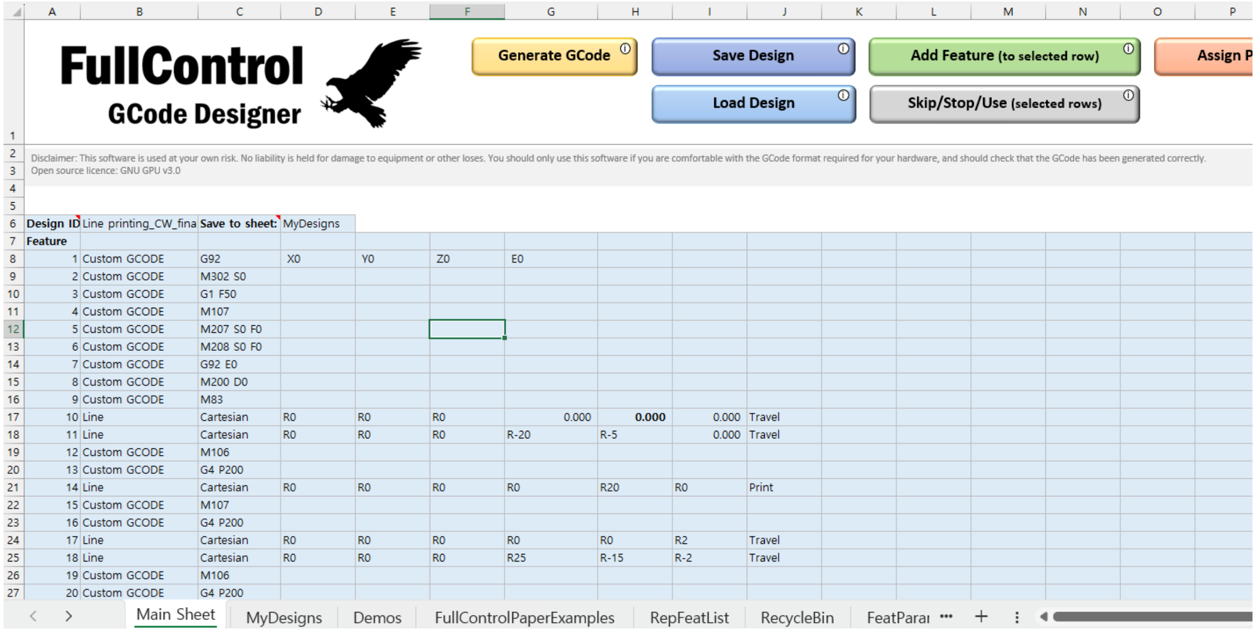


**Fig. S2**. User interface of FullControl software.

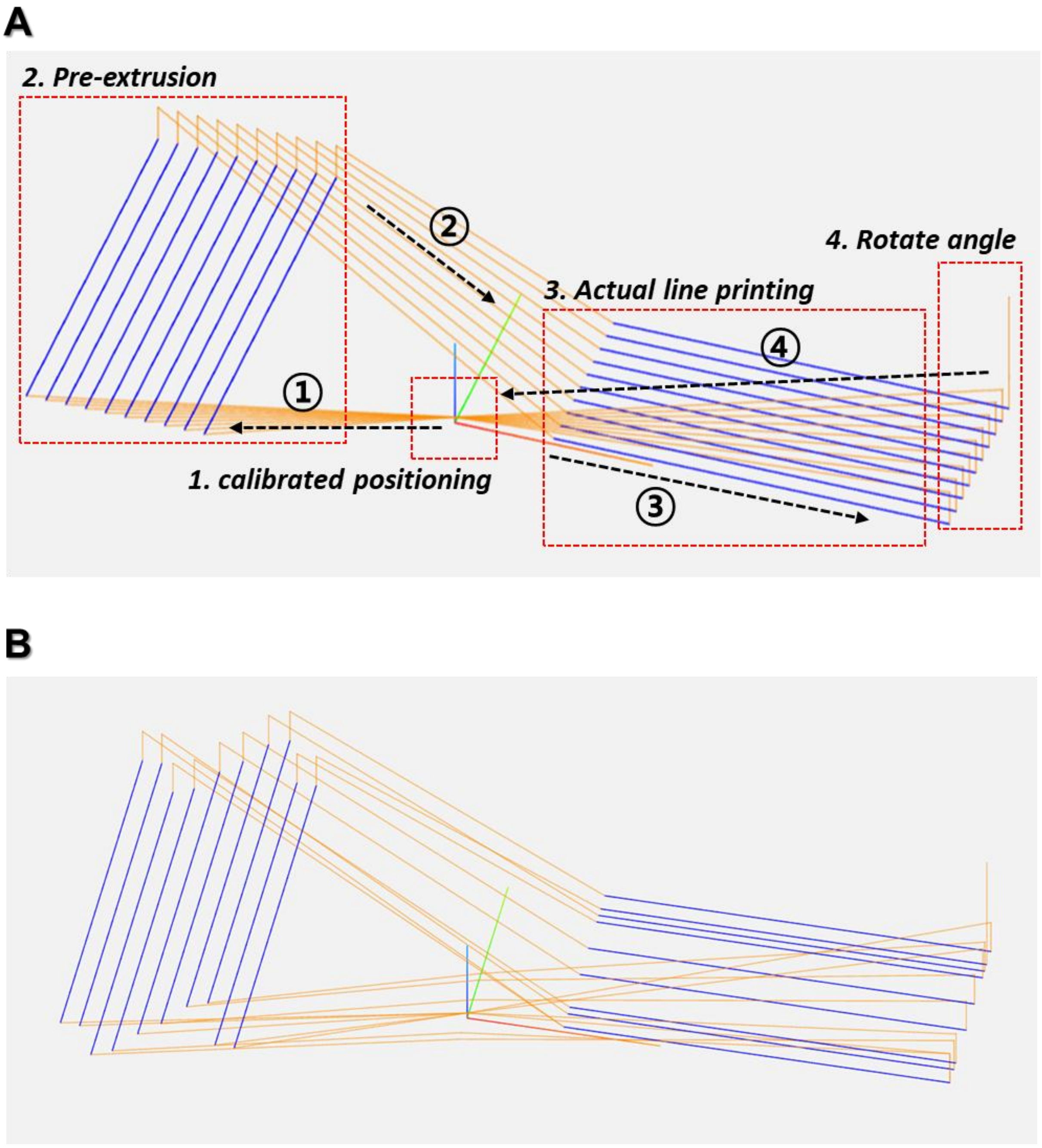


**Fig. S3**. G-code paths: A) original G-code path with the printing procedure and B) G-code path after applying the calibrated results.

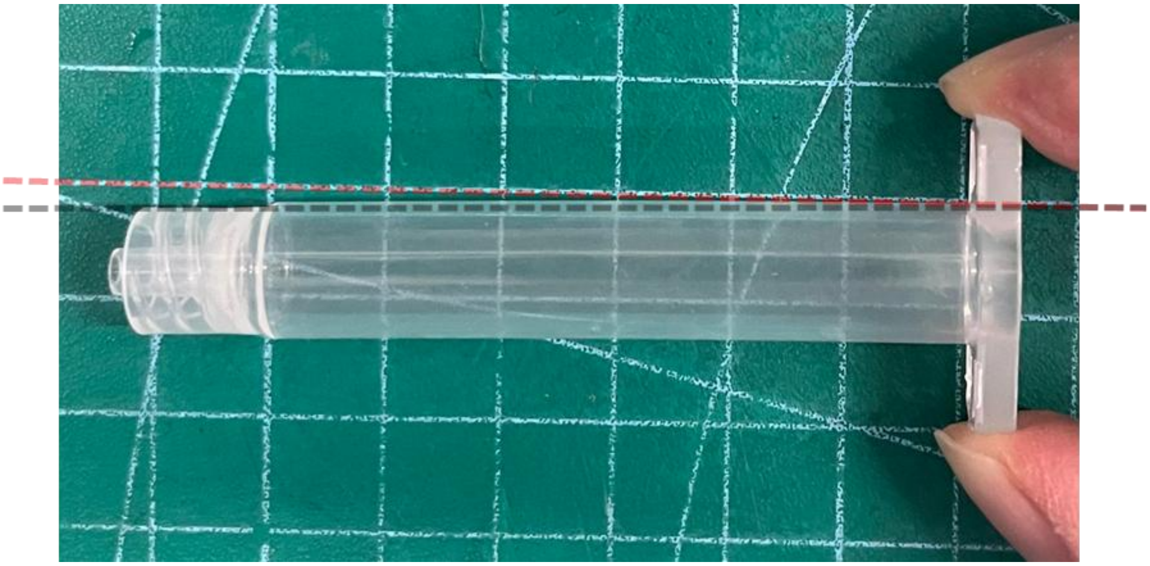

**Fig. S4**. Bent plastic syringe cartridge (red line: straight reference line; black line: syringe bending).

**Table S1**. Calibrated results for the X-, Y-, and Z-axes.

| Nozzle | N1 | N2 | N3 | N4 | N5 | N6 | N7 | N8 | N9 | N10 |
|---|---|---|---|---|---|---|---|---|---|---|
| X-axis (CW) | E3.590 | E3.660 | E3.510 | E3.635 | E3.615 | E3.520 | E3.570 | E3.570 | E3.580 | E3.560 |
| X-axis (CCW) | E-3.585 | E-3.625 | E-3.505 | E-3.640 | E-3.620 | E3.525 | E-3.595 | E-3.570 | E-3.560 | E-3.560 |
| Y-axis | 0.0 | -2.5 | 3.0 | 1.5 | 1.0 | 1.0 | 0.0 | 3.5 | 0.0 | 0.0 |
| Z-axis | 0.0 | 0.0 | 0.0 | 0.0 | 0.1 | 0.2 | 0.1 | 0.1 | 0.0 | 0.1 |

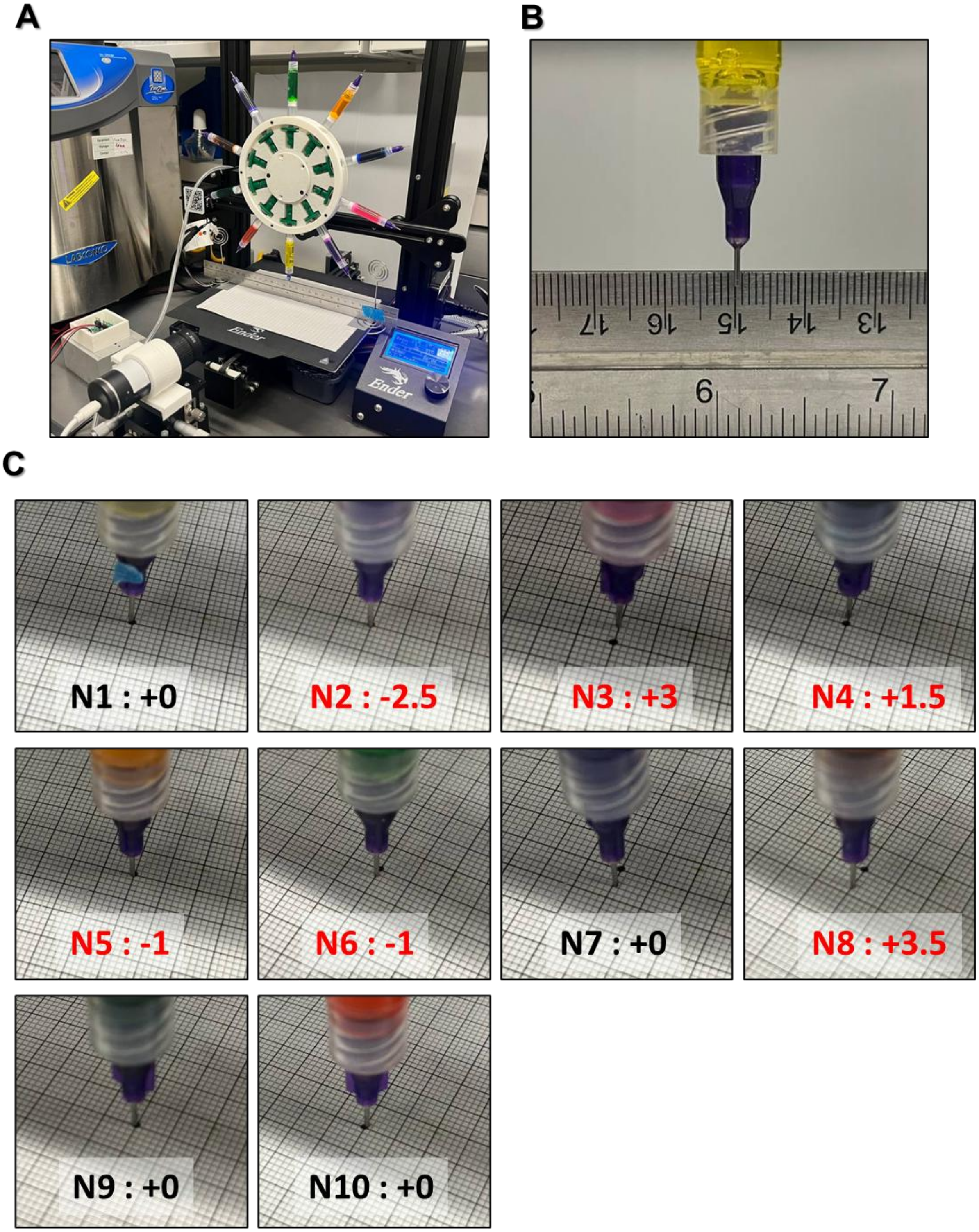


**Fig. S5**. Manual calibration process: A) MNB system setup for calibration with a camera and ruler, B) captured image from the camera, and C) Y-axis calibration results measured on a grid sheet.

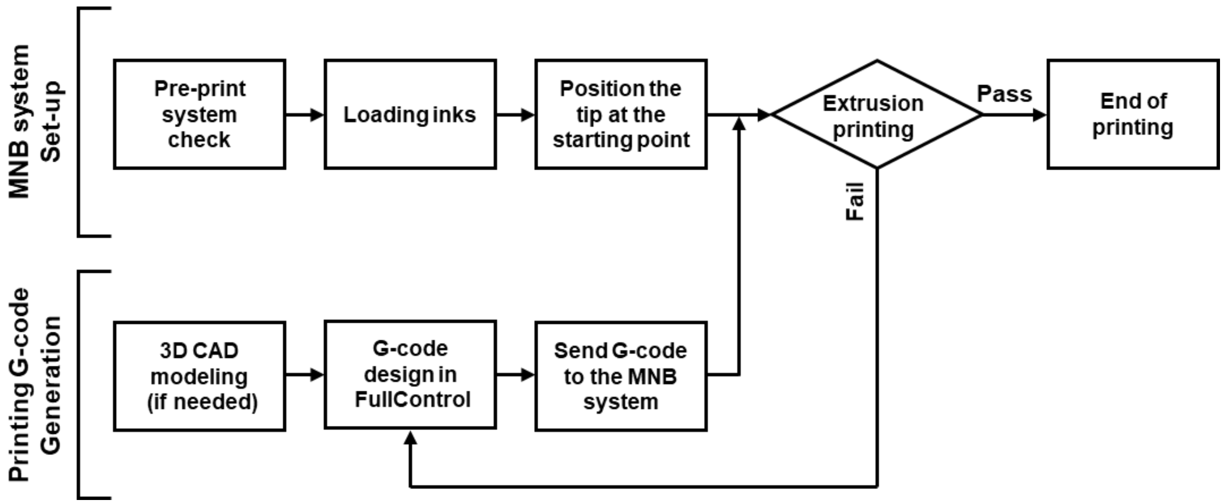


**Fig. S6**. Flowchart of printing process for the G-code generation and printing

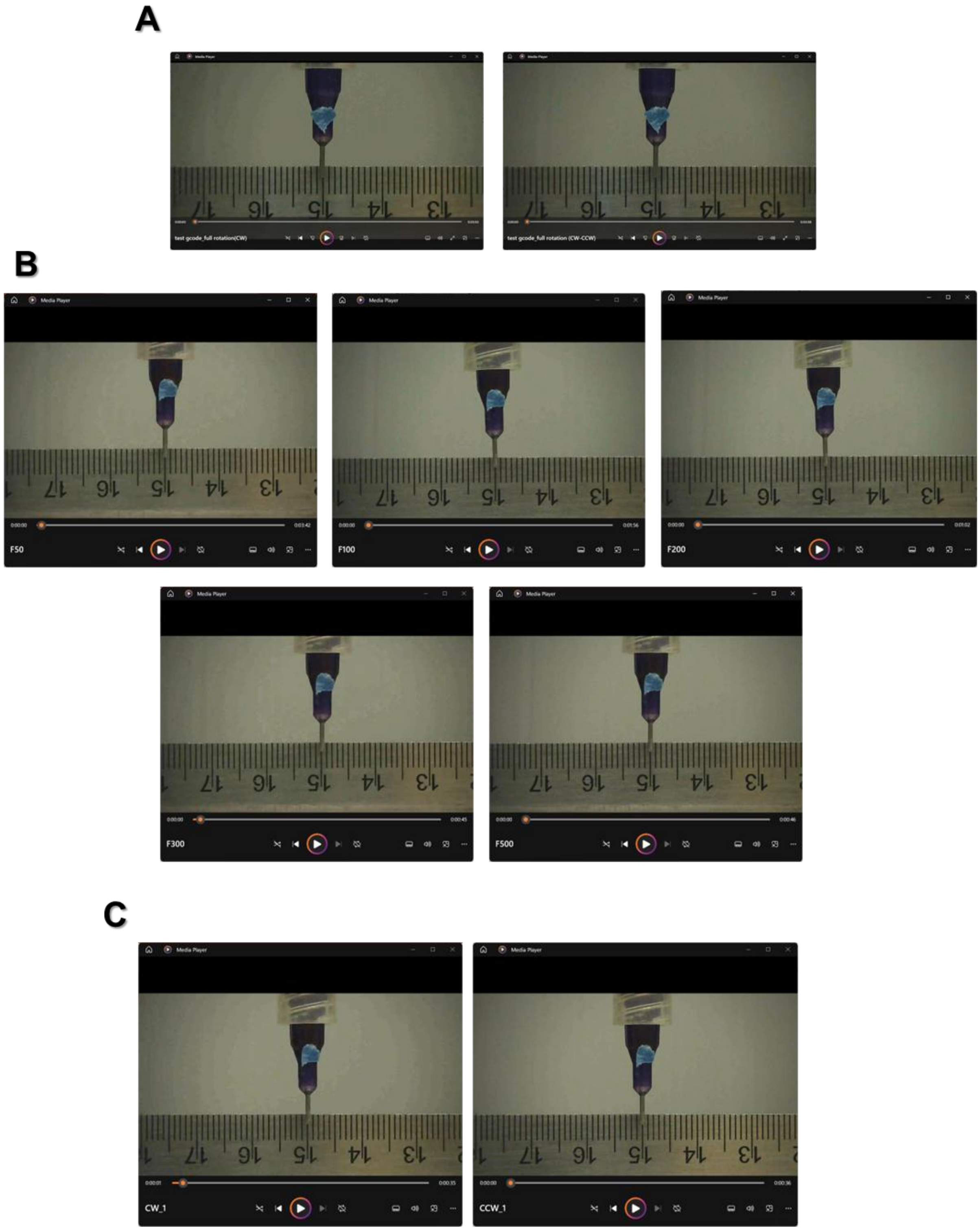


**Fig. S7**. Representative images extracted from videos for calibration: A) repeatability test, B) speed calibration, and C) single-nozzle calibration.

**Supplementary Video_S1_calibration**

**Supplementary Video_S2_line printing**

**Supplementary Video_S3_auxetic scaffold**
